# Taxonomic and metabolic diversity of Actinobacteria isolated from faeces of a 28,000-year-old mammoth

**DOI:** 10.1101/2022.12.22.521380

**Authors:** Doris A. van Bergeijk, Hannah E. Augustijn, Somayah S. Elsayed, Joost Willemse, Victor J. Carrión, Mia Urem, Lena V. Grigoreva, Maksim Y. Cheprasov, Semyon Grigoriev, Bas Wintermans, Andries E. Budding, Herman P. Spaink, Marnix H. Medema, Gilles P. van Wezel

## Abstract

Ancient microbial communities of permafrost soils and frozen animal remains represent an archive that has barely been explored. This yet unexplored microbial world is a vast resource that can provide us with new evolutionary insights, metabolic pathways and novel chemistry. Here, we reveal that Actinobacteria isolated from a faecal sample from the intestinal tract of a 28,000-year-old Siberian mammoth are phylogenetically and metabolically distinct from currently known modern siblings. Ancient *Micromonospora, Oerskovia, Saccharopolyspora, Sanguibacter* and *Streptomyces* species were successfully revived and their genome sequences resolved. Surprisingly, the genomes of the ancestors show a large phylogenetic distance to strains isolated today and harbour many novel biosynthetic gene clusters that may well represent uncharacterised biosynthetic potential. Metabolic profiles of the strains display production of known molecules like antimycin, conglobatin and macrotetrolides, but the majority of the mass features could not be dereplicated. Our work provides a snapshot into Actinobacteria of the past, yielding unexplored genomic information that is not yet present in current databases.

## Introduction

Bacteria are key to life on Earth; they are major players in biogeochemical nitrogen cycling ^1^, decompose organic matter ^2^, and provide eukaryotic hosts with essential nutrients and protection against biotic and abiotic stress ^3, 4^. Bacteria also produce a wide range of natural products, among them many have applications in medicine, biotechnology and agriculture ^5, 6^. It is predicted that we have only uncovered a small percentage of the microbial world; the majority of microorganisms resist cultivation and many bacterial taxa have hardly been explored ^7^. Indeed, scientists have so far primarily accessed easily accessible environments, while numerous bacteria exist in remote niches such as deep-sea sediments, caves, and permafrost soils ^8, 9^. This uncharacterized microbiology likely represents an important reservoir of biological information and may among others be harnessed for drug discovery.

Insights into microbial communities in ancient environments provide us with a glimpse into the past and may teach us important lessons on the evolution of microbial and chemical diversification. Isolation and metagenome sequencing of ancient bacteria has provided important knowledge on bacterial evolution, changes in microbiome composition, ancient diseases and potential chemical novelty ^10-13^. For example, large-scale *de novo* assembly of microbial genomes from human paleofaeces samples revealed previously undescribed gut microorganisms and a markedly higher abundance of mobile genetic elements in our ancestral gut microbiome compared to modern industrial gut microbiomes ^10^. A metagenomic survey of ancient Alaskan soil confirmed that homologues of different resistance genes existed in ancient bacteria, showing that antibiotic resistance predates the modern selective pressure of clinical antibiotic use ^11^. Furthermore, ancient Actinobacteria isolated from thousands-year-old Arctic and Antarctic sediments showed promising bioactivity against drug-resistant pathogens, and genome mining showed low similarity to known antibiotics ^12, 13^. These studies illustrate how ancient sources can lead to evolutionary insights, as well as novel micro-organisms and chemistry.

The New Siberian Islands, located between the Laptev Sea and the East Siberian Sea, contain permafrost deposits that have been preserved for more than 200,000 years ^14^. These islands are considered a time capsule to ancient biology and have been an important source of ancient animal remains, including mammoths ^15^. In August 2012, an adult female woolly mammoth (*Mammuthus primigenius*) was recovered on Maly Lyakhovsky Island (74°07′ N, 140°40′ E) ^16, 17^. The mammoth carcass was submerged in permafrost, exposing skull, post-cranial elements and partial trunk. The lower part of the body was surrounded by almost pure ice and included lower parts of the head, distal portion of the trunk, chest, abdomen, front legs and distal half of the right hind leg ^16-18^. The mammoth was determined to be about 28,500 years old and the remains contained exceptionally well-preserved soft tissues. The skin had retained its elasticity and mummification of the carcass was minimal ^16-18^. In February 2014, the pristine specimen was transported to Yakutsk for investigation.

The finding of this exceptionally well-conserved ancient specimen provided a unique opportunity to explore its ancient microbiome. Its well-preserved soft tissues included the intestinal tract, which was used to extract a faecal sample for isolation of bacteria. We focused specifically on members of the highly diverse phylum Actinobacteria, well known for their capability to produce an unprecedented diversity of specialised metabolites. These bacteria evolved ∼2,700 million years ago and reproduce as exospores, which should allow their long-term survival in permafrost ^19^. Their genomes typically contain many biosynthetic gene clusters (BGCs) that encode the cellular machinery for the biosynthesis of natural products. Although many bioactive compounds produced by Actinobacteria have been identified, genomic research shows that a large part of the metabolic diversity of this phylum is still unexploited ^20^. In this study we isolated Actinobacteria from this extraordinary ancient sample, compared the genomes to those of their closest modern-day neighbours and analysed their bioactive potential. Sequencing of the six isolated Actinobacteria revealed significant phylogenetic distance to currently known strains, with yet uncharacterised biosynthetic potential.

## Results

### Isolation of Actinobacteria from mammoth faeces

The remarkable diversity of Actinobacteria and their specialised metabolites is the result of millions of years of evolution ^21, 22^. However, the evolutionary drivers that have shaped this metabolic diversity remain largely unknown. Isolation of Actinobacteria from ancient samples may allow a glimpse into the past and thus provide insights on how natural product biosynthesis evolved over this period of time. The discovery of a well-preserved mammoth on Maly Lyakhovsky Island ^16, 17^ (Fig. S1) provided a unique opportunity to recover ancient Actinobacteria and compare their genomes and biosynthetic potential to their modern descendants. From March 10-14 2014, we joined an international team of researchers during the dissection of this extraordinary specimen. During the dissection, every day a deeper layer of the mammoth tissue became accessible, as the specimen gradually thawed. When the thawing had proceeded far enough to explore the abdominal cavity, we found a large part of the intestines fully intact, with the omentum still attached. To extract faecal samples, the intestines were exposed and a 60 cm intestinal specimen was gathered from the remains (Fig. S1). Caution was taken to carefully remove the faecal sample, and the specific sample we eventually used for isolation of the Actinobacteria was then extracted from the very core of the large faecal sample, so as to reduce the risk of contamination to the absolute minimum. The intestinal lumen was thoroughly inspected for defects, and faeces was taken and stored using sterile materials. Two perpendicular incisions were made and the intersection was folded over, exposing the intestinal lumen. Faecal samples were carefully taken from the lumen under sterile conditions, thereby avoiding any cross-contamination (Fig. S1).

A fraction of the collected faeces sample was homogenised in sterile dH_2_O and plated onto various media selective for Actinobacteria ^23^. Plates were incubated aerobically at 4 °C and 30 °C, and anaerobically at room temperature. We were able to recover Actinobacteria from the 28,500 year sample. Strains were selected based on their filamentous morphology and grown on different media for phenotypic discrimination, resulting in the isolation of six morphologically distinct strains. The majority of the strains were isolated using selective humic acid agar and/or glucose agar plates incubated at 30 °C. No filamentous bacteria were observed after anaerobic incubation.

### Taxonomic and phenotypic profiling of the Actinobacteria

To determine the taxonomic origin of the isolates and gain insights into the relatedness to current known bacterial species, total genomic DNA was isolated from each of the isolates and full genome sequences were obtained using a combination of Nanopore and Illumina sequencing. Comparison of the 16S rRNA sequence to those within the EzBioCloud database revealed that the isolates belonged to five genera of Actinobacteria, namely *Sanguibacter* (M9), *Micromonospora* (M12), *Oerskovia* (M15), *Saccharopolyspora* (M46) and *Streptomyces* (M10 and M19). To obtain a more detailed taxonomic classification, a maximum-likelihood tree was constructed based on the genome sequences of the six isolates and that of 578 Actinobacteria representing six bacterial families and over 40 different genera (Fig. 1). The relatedness of the isolates and neighbouring organisms was determined by calculating the average nucleotide identity (ANI) score in an all-to-all genome comparison. The ANI score revealed a surprisingly large phylogenetic distance between the mammoth isolates and currently known strains, with similarity scores ranging from 82-89% (Table S1). After all, while 28,500 years is a long period of time, on the evolutionary scale it is short, especially considering that streptomycetes evolved about 2,700 million years ago. Of the isolates, the largest phylogenetic distance was found for *Streptomyces* sp. M19 and its closest neighbours with an ANI score <83%. The species delineation threshold typically lies at approximately 95% gene identity ^24^, which suggests that the isolates may be novel ancient species.

**Figure 1.**
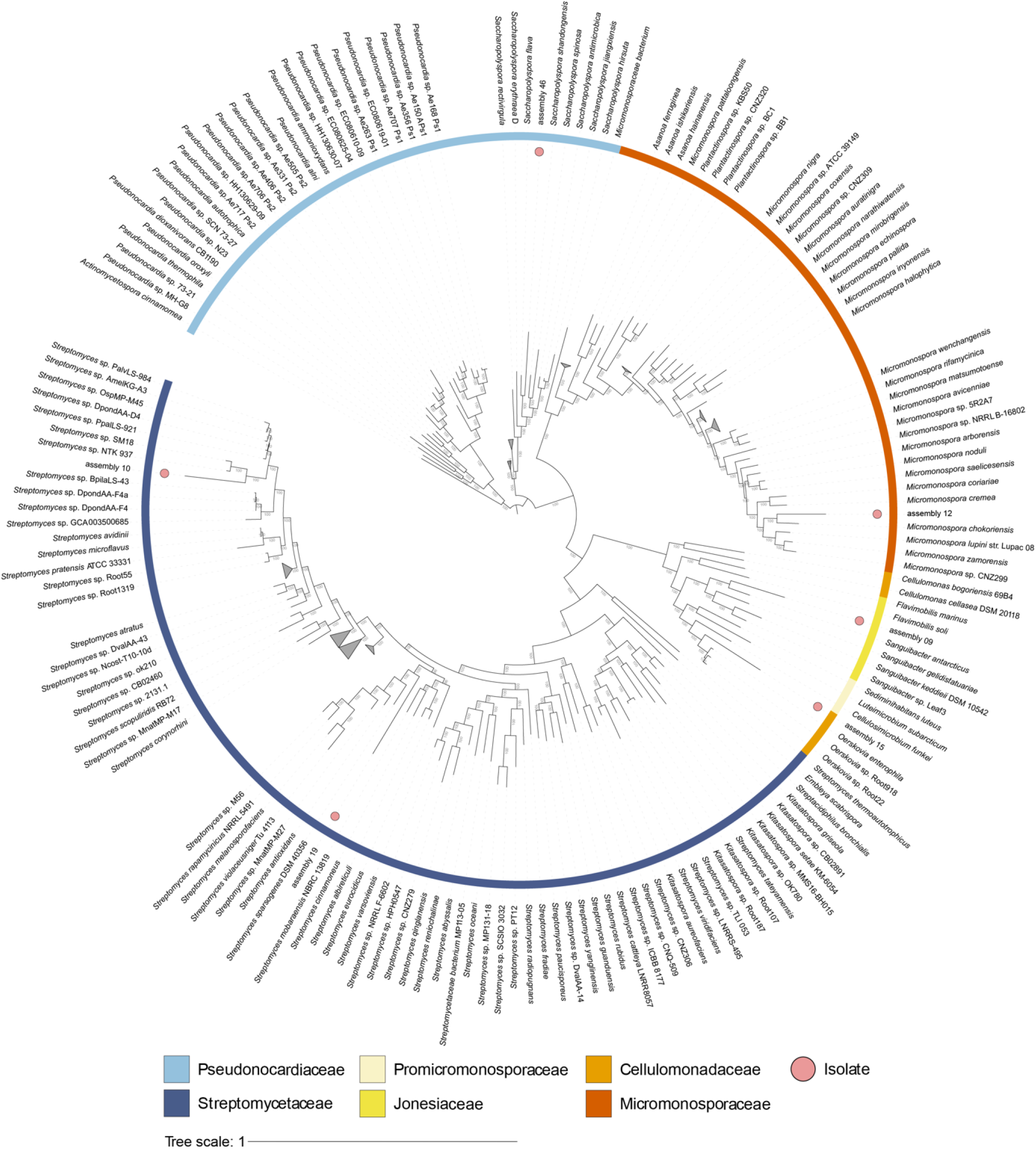
Phylogeny of mammoth isolates and their closest known neighbours. Maximum-likelihood tree of the six isolates compared to 578 Actinobacteria of six bacterial families (colour-coded in the outer ring), subdivided into 40 genera. The tree was rooted using *Pseudonocardia* sp. as outgroup and the numbers on the tree branches represent the bootstrap values in percentages of a total of 1000 bootstraps. Circles indicate the location of the novel isolates. The phylogenetic analysis suggests the following likely classification based on their nearest neighbours: M9, *Sanguibacter*; M12, *Micromonospora*; M15, *Oerskovia*; M10 & M19, *Streptomyces*; M46, *Saccharopolyspora*.

Taxonomic analysis revealed a significant phylogenetic difference between the genomes of the isolates and those of their closest known neighbours, as indicated by the branch lengths (Fig. 1). This was especially surprising for the *Streptomyces* isolates, which were compared to more than 300 fully annotated genomes. We wondered whether the low similarity was a result of the ancient origin of the strains or whether this was related to the underexplored environment the strains were sampled from. Therefore, we analysed the phylogenetic distance between other strains isolated from extreme environments and their closest neighbours. For this we selected the deep sea isolates *Streptomyces* sp. NTK 937 ^25^ and *Streptomyces* sp. SCSIO 3032 ^26^, the desert isolate *Streptomyces thermolilacinus* SPC6 ^27^, and *Streptomyces* sp. BF-3 and *Streptomyces* sp. 4F isolated from the Great Salt Plains in Oklahoma ^28^ (Table S2). In general, the results show higher relatedness to the closest neighbours of these strains (ANI scores > 93%, compared to 82-89% for our isolates), with the exception of *Streptomyces* sp. SCSIO 3032 (ANI: 85.48%) and *Streptomyces thermolilacinus* SPC6 (ANI: 88.22%). It should be noted that the closest known neighbour of *Streptomyces thermolilacinus* SPC6 is an isolate from island soil, while this strain itself was isolated from a soil sample collected in the desert. Additionally, *Streptomyces* sp. SCSIO 3032 and its neighbour *Streptomyces* sp. MP131-18 both originate from deep-sea samples, but from different sampling locations. Thus, we cannot exclude that the low level of similarity observed between the genomes of our strains and publicly available genomes may be due to the lack of bacteria isolated from similar environments, rather than the age of the strains themselves. Further analyses of ancient samples should shed more light onto this important question.

### Morphological characterization of the isolates

To obtain a more detailed view of the morphology of the Actinobacterial colonies, the isolates were grown on SFM agar and subsequently analysed by microscopy. Stereomicroscopic analysis revealed a wide range of phenotypes (Fig. 2A). *Sanguibacter* sp. M9 produced bright yellow round colonies, *Streptomyces* sp. M10 produced cream-coloured substrate mycelia, a grey aerial spore mass, and a dark diffusible pigment, *Micromonospora* sp. M12 produced orange-coloured folded colonies, *Oerskovia* sp. M15 produced white colonies, and *Saccharopolyspora* sp. M46 produced cream-coloured folded colonies. *Streptomyces* sp. M19 displayed a heterogeneous phenotype. When this isolate was grown on SFM agar, two colony phenotypes were observed: a fully developed phenotype, and a variant with strong yellow pigmentation, sparse aerial mycelia and lack of spores. Morphological heterogeneity was also observed within single colonies (Fig. S2), suggesting a high tendency to genetic heterogeneity ^29^. Sequencing of the 16S rRNA strongly suggests that all morphological variants were indeed phenotypes of the same strain.

**Figure 2.**
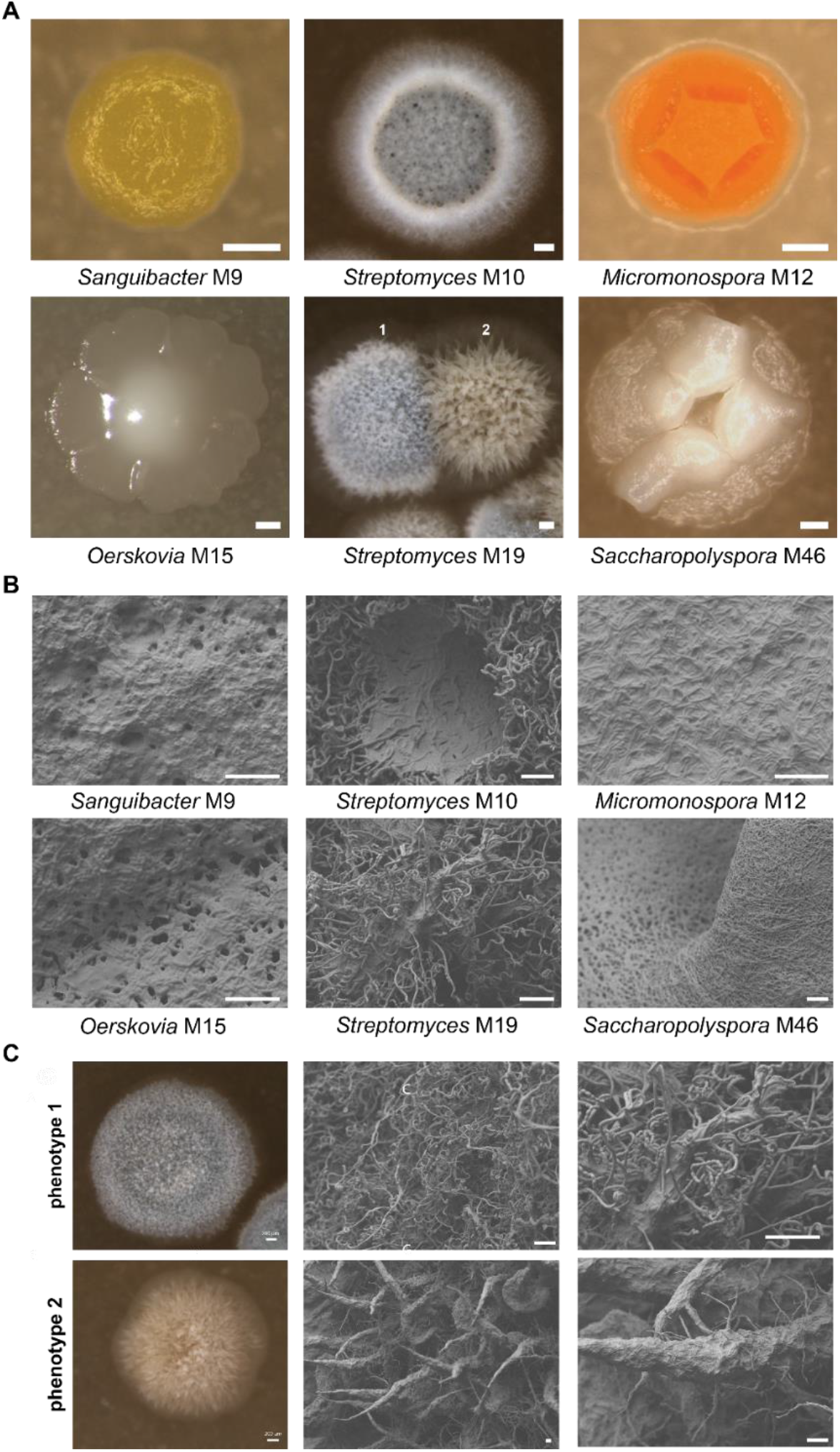
Phenotypic characterization of Actinobacteria isolated from a faecal sample of a 28,000-year-old mammoth. Strains were grown on SFM agar plates for 9 days. **(A)** Stereomicrographs of isolates *Sanguibacter* sp. M9, *Streptomyces* sp. M10, *Micromonospora* sp. M12, *Oerskovia* sp. M15, *Streptomyces* sp. M19, and *Saccharopolyspora* sp. M46 (for phylogenetic analysis see Fig. 1). *Streptomyces* sp. M19 showed two distinct phenotypes: fully developed colonies (1) and colonies with a bald appearance (2). Scale bar: 200 µm. **(B)** Scanning electron micrographs of isolates M9, M10, M12, M15, M19, and M46. Scale bar: 10 µm. **(C)** Scanning electron micrographs showing significant differences in morphology between the two colony phenotypes observed when *Streptomyces* sp. M19 is grown on SFM. Images of phenotype 1 show spirals of spore chains. Images of phenotype 2 show large spikes made up of hyphae and extracellular matrix. Also in the fully developed colony, such spikes can be found but in low frequency.16S rRNA sequencing strongly suggests that these morphological variants are phenotypes of *Streptomyces* sp. M19. Scale bar: 10 µm.

To obtain more insights into the morphology of the strains at high resolution, colonies were subjected to Scanning Electron Microscopy (SEM) (Fig. 2B). Colonies of *Streptomyces* sp. M10 produced hairy spores, whereby the aerial mycelium consisted of both smooth and hairy hyphae, with dark-pigmented droplets on top of the colonies. *Saccharopolyspora* sp. M46 colonies consisted of a thick layer of interwoven mycelium made up of hyphae and extracellular matrix. SEM studies of the two distinct phenotypes of *Streptomyces* sp. M19 revealed spiral spore chains in one variant (phenotype 1), while we failed to identify spores in the other (phenotype 2) (Fig. 2C). Instead, the non-sporulating colonies produced large spikes made up of hyphae embedded in an extracellular matrix. These spikes could also be found in the fully developed colony, but in low abundance. Colonies of isolates *Sanguibacter* sp. M9, *Micromonospora* sp. M12, and *Oerskovia* sp. M15 were covered by an extracellular matrix and could therefore not be further characterised by SEM.

Taken together, the taxonomic and phenotypic analyses show that we have successfully isolated a diverse range of Actinobacteria, whereby *Streptomyces* sp. M19 stood out with a highly heterogeneous phenotype and low similarity to its closest neighbours. Moreover, the low ANI scores suggest that all isolates represent novel species, yielding unexplored genomic information that is not yet present in current databases.

### Biosynthetic potential of the Actinobacteria

We then studied the biosynthetic gene clusters (BGCs) of the isolates, to assess their relatedness to BGCs that are currently available in the databases. For this, the genome sequences were analysed using antiSMASH ^30^. This identified a total of 179 putative BGCs, namely four in *Oerskovia* sp. M15, six in *Sanguibacter* sp. M9, 19 in *Saccharopolyspora* sp. M46, 23 in *Micromonospora* sp. M12, 31 in *Streptomyces* sp. M10 and 34 in *Streptomyces* sp. M19 (Fig. 3A). Over 70% of the total BGCs shared less than 50% KnownClusterBlast similarity to BGCs within the MiBIG database, with 23% showing no significant similarity to any known BGC. This amounts to a similar amount of biosynthetic novelty compared to many genomes from contemporary strains (Table S3). The genomes contained relatively high numbers of genes encoding enzymes associated with the biosynthesis of terpenes, in particular for *Micromonospora* sp. M12 (five), *Streptomyces* sp. M19 (four), *Streptomyces* sp. M10 (seven) and *Saccharopolyspora* sp. M46 (six).

**Figure 3.**
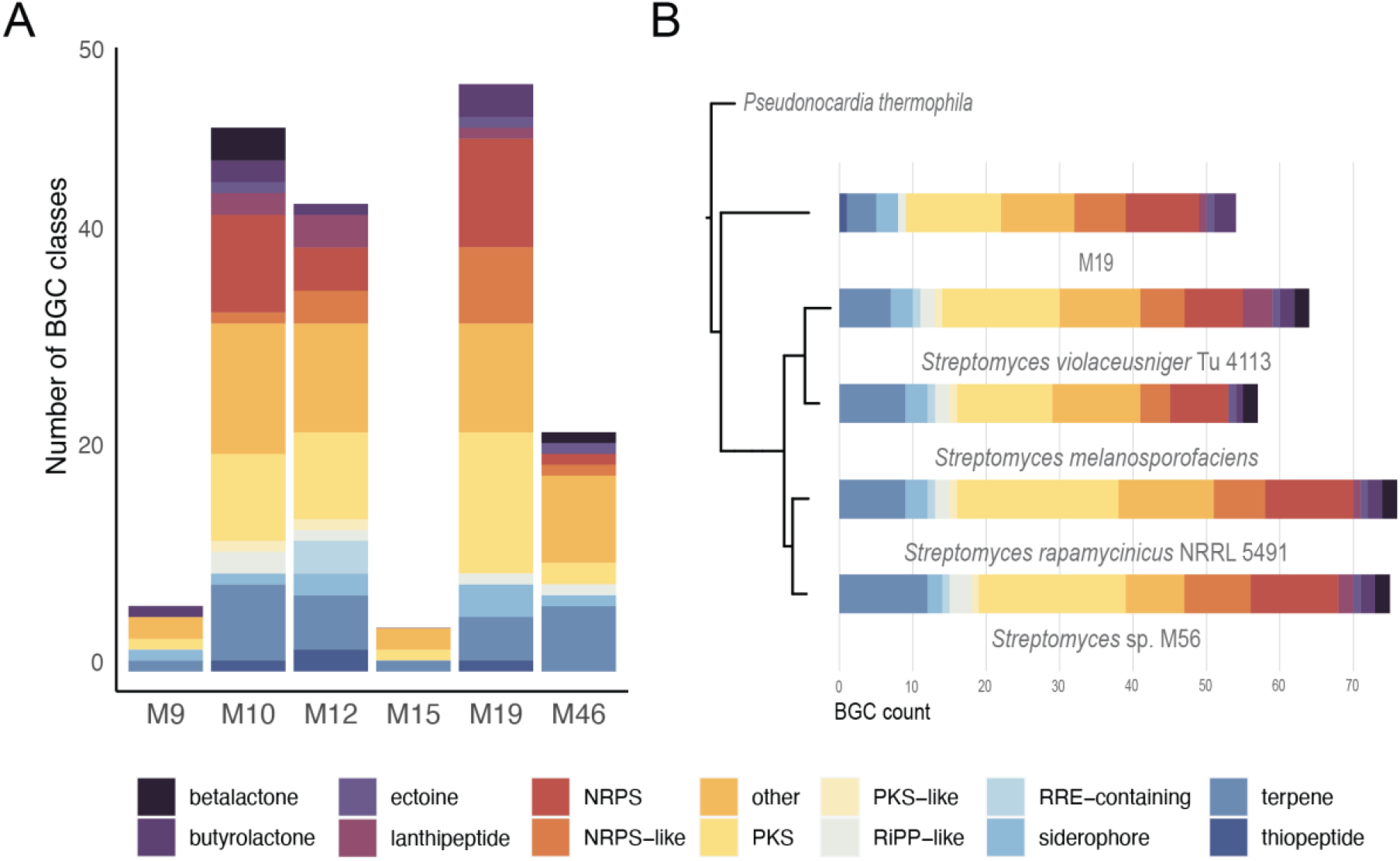
Classification of BGCs predicted in the actinobacterial genomes. **A)** BGC classes for each isolate predicted with antiSMASH v6.0. Known types of BGCs representing <1% of all BGCs were grouped into the “other” category. The large number of BGCs in *Streptomyces* spp. M10 and M19, *Micromonospora* sp. M12 and *Saccharopolyspora* sp. M46, is particularly noteworthy. **B)** Overview of all predicted BGC classes for isolate M19 and its closest neighbours *Streptomyces violaceusniger* Tu 4113, *Streptomyces rapamycinicus* NRRL 5491, *Streptomyces* sp. M56 and *Streptomyces melanosporofaciens*. The comparison shows that only two BGCs are shared between all three isolates, namely BGCs for desferrioxamines and for ectoine. Moreover, the genome of *Streptomyces* sp. M19 contains a greater variety of BGC classes, with a surprisingly high proportion of NRPS BGCs.

The genome sequence of *Streptomyces* sp. M19 showed the largest distance in terms of ANI scores to its nearest neighbours. We therefore subjected the strain to detailed analysis of its biosynthetic potential, to obtain insights into the chemical diversity of its specialised metabolites. *Streptomyces* sp. M19 is predicted to have seven closest neighbours (Table S2). Out of these seven, we selected four strains with the highest quality assemblies for a comparative genome analysis, namely *Streptomyces violaceusniger* Tu 4113, *Streptomyces rapamycinicus* NRRL 5491, *Streptomyces* sp. M56, *Streptomyces melanosporofaciens* (Fig. 3B). The M19 genome had a slightly lower number of predicted BGCs (34) as these nearest neighbours (47-54 BGCs). Surprisingly, *Streptomyces* sp. M19 only shares two BGCs with its nearest neighbours, namely for desferrioxamines and for ectoine. The M19 genome encodes a high percentage of non-ribosomal peptide synthetases (NRPSs) and polyketide synthases (PKSs), and also has BGCs for butyrolactones, unusual polyketides / fatty acids (linked to a heterocyst glycolipid synthase-like PKS, and another unusual type of ketosynthase related to those found in ladderane lipid biosynthetic pathways) a likely lasso peptide, an aryl polyene, and an aminoglycoside/aminocyclitol; none of these natural product classes could be found in modern nearest neighbours. Comparative genomic analysis of *Streptomyces* sp. M19 and its four neighbours displays the low similarity of approximately 82% and only few corresponding BGCs (Fig. S3).

Additionally, metagenome reads from mammoth faeces and ice surrounding the mammoth were obtained to further exclude the possibility of contamination. The raw reads were mapped against the complete genomes and 16S regions of the six isolates and normalized for sequencing depth. Resulting read mapping numbers show only sparse mapping to our isolates in both the faecal and ice sample (Fig. S4). To distinguish between reads mapping in conserved or highly specific regions in the genome, we performed a read mapping analysis against the predicted BGCs. This showed no mapping to core biosynthetic genes in the ice sample. In the faecal sample, we could observe one read mapping to a redox-cofactor cluster of *Streptomyces* sp. M19 (coverage of 3.48%) and one to a nystatin cluster of *Saccharopolyspora* sp. M46 (coverage of 0.13%).

### Antibiotic activity and bioactive metabolites produced by the mammoth isolates

Next, the antibiotic-producing potential of the strains was assessed under six different culturing conditions. These were four agar-based media, namely the nutrient-rich Nutrient Agar (NA) and International *Streptomyces* Project 2 (ISP2) and the nutrient-poor Czapek Dox and Minimal Media supplemented with mannitol and glycerol (MM), and the liquid media versions of ISP2 and MM. We used the Gram-positive *Bacillus subtilis* 168 and the Gram-negative *Escherichia coli* ASD19 and *Pseudomonas aeruginosa* PA01 as indicator strains. Different types of growth inhibition were observed, namely complete inhibition, strong reduction in number of colony-forming units (cfu), and impaired growth (Fig. 4). Under the conditions tested, the isolates displayed the strongest antibacterial activity when grown on NA. Therefore, further metabolic analysis was done on samples isolated from NA-grown cultures.

**Figure 4.**
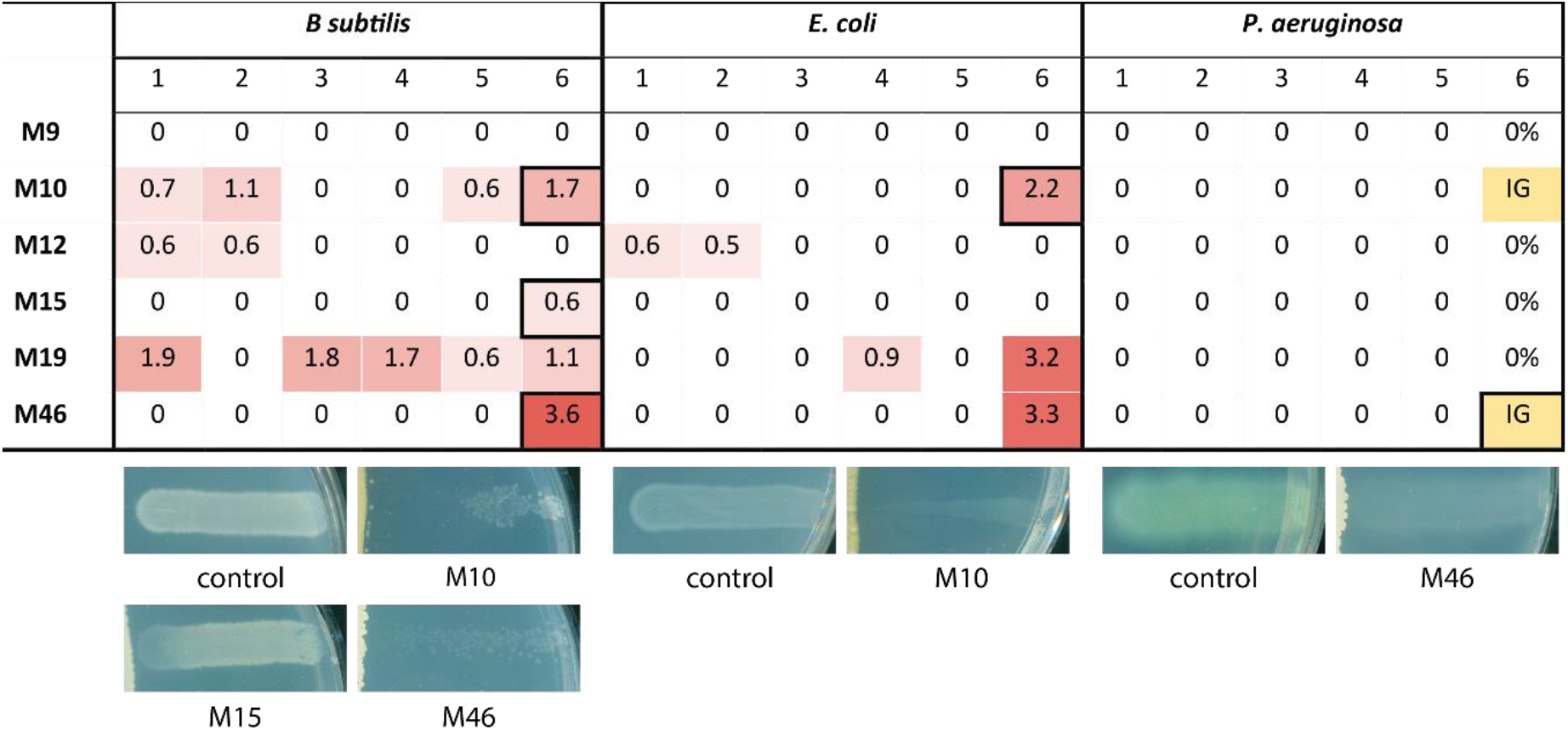
Antimicrobial activity of the actinobacterial isolates. After 7 days of growth, the antimicrobial activity of the mammoth isolates was assessed against different indicator strains using different growth media and methods: 1 = MM, soft agar overlay; 2 = Czapek Dox, soft agar overlay; 3 = liquid culture ISP2; 4 = liquid culture MM; 5 = ISP2, cross streak, 6 = NA, cross streak. The zone of inhibition (cm) is indicated for each isolate and related to a colour scale (*n* = 3). Most activity was observed in the cross-streak assay on NA. The fields selected with a black border refer to the examples displayed below the table that illustrate the different types of growth inhibition observed. IG, impaired growth.

Besides soluble natural products, also volatile organic compounds (VOCs) may have antibacterial activity ^31, 32^. To investigate whether (some of) the antimicrobial activity may have been due to the production of antibacterial VOCs, the isolates were grown on plates where the Actinobacteria were separated from the indicator strains by an impermeable polystyrene divider. Interestingly, growth of *E. coli* was completely inhibited by VOCs produced by isolates *Streptomyces* sp. M10, *Streptomyces* sp. M19, and *Saccharopolyspora* sp. M46 (Fig. S5). While *Micromonospora* sp. M12 did not display antibacterial activity in the bioactivity assay described above, it partially inhibited growth of *E. coli* in the volatile assay. None of the strains produced VOCs that could inhibit growth of *B. subtilis*. These data indicate that the observed antibacterial activity of strains M10, M19, and M46 against *E. coli* was at least in part caused by VOCs, while the bioactivity of isolates M10, M15, M19, M46 against *B. subtilis* was solely caused by the production of soluble antibiotics.

To gain more insights into the soluble antibiotics produced by isolates *Oerskovia* sp. M15, *Streptomyces* sp. M10, *Streptomyces* sp. M19 and *Saccharopolyspora* sp. M46, the strains were streaked on NA plates and grown for seven days. Metabolites were extracted using ethyl acetate (EtOAc) and tested for bioactivity against the different indicator strains. The crude extracts of *Streptomyces* sp. M10 and M19 showed activity against *B. subtilis*, while *E. coli* and *P. aeruginosa* were not inhibited (Fig. S6). MS/MS data were analysed using Global Natural Products Social molecular networking (GNPS) ^33^, resulting in a molecular network containing 2886 nodes clustered in 223 spectral families (Fig. 5). The highest number of unique nodes (491) were attributed to isolate M10, while the lowest number (280) was attributed to isolate M46. 44 nodes were unique to *Streptomyces* isolates M10 and M19. Dereplication based on matching MS/MS spectra against the GNPS spectral library annotated several *m/z* values as being known bioactive metabolites: antimycin A1 (**1**), antimycin A2 (**2**), and the macrotetrolide monactin (**3**) and related bonactin (**4**) and homononactyl homononactate (**5**) in the extracts of M10, and conglobatin (**6**) in the extracts of M19 (Fig. 5). To confirm these findings, we analysed the KnownClusterBlast output of antiSMASH for presence of the responsible BGCs. In the genome of M10, antiSMASH identified an antimycin BGC and a BGC with 75% similarity to a macrotetrolide BGC (MIBiG cluster BGC0000243). The macrotetrolide-associated BGC from strain M10 lacks three genes compared to the reference BGC, two encoding hypothetical proteins and one encoding an inositol monophosphatase-like enzyme. M19 harbours a BGC with low (36%) similarity to the conglobatin BGC (Fig. S7). Although the overall match for this cluster is lower, we could detect homologues of all known core conglobatin genes (*congA-E*) with the same domain architecture as in the reference BGC. Taken together, the isolates produced some known bioactive molecules, but the majority of the mass features could not be dereplicated, suggesting the production of novel chemistry.

**Figure 5.**
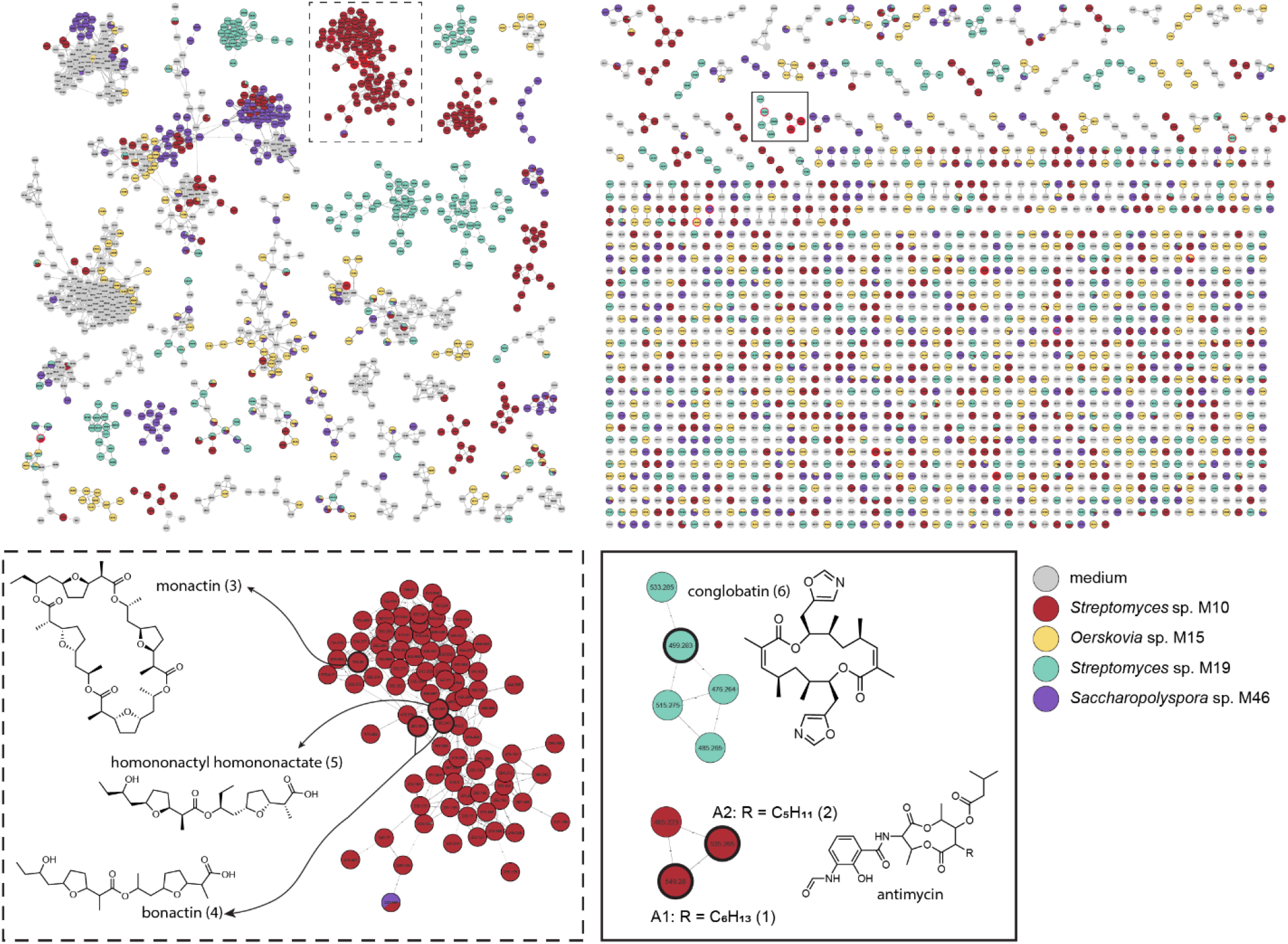
Molecular network of the ions detected in crude extracts and dereplication of bioactive compounds. A pie chart was mapped to the nodes which represents the abundance of each *m/z* value in the crude extracts of the different bioactive isolates (M10, M15, M19 and M46) and the medium blank. Nodes highlighted represent dereplicated known bioactive metabolites. The isolates were grown for seven days on NA agar plates (*n* = 3).

## Discussion

A huge proportion of the bacterial world remains uncharacterised, representing an important reservoir of biological information and new chemistry as the possible basis for future drugs ^7, 8, 34^. Metagenome sequencing, varying culturing techniques, and isolation of bacteria from rare environments is gradually revealing part of this microbial dark matter, and ancient samples are a promising and underexplored resource ^10, 11^. We had the unique opportunity to sample faeces from the intestinal tract of an exceptionally well-preserved 28,000-year-old mammoth, which we used to isolate ancient Actinobacteria ^16^. These Actinobacteria belonged to the well-studied genus *Streptomyces* and to the underexplored genera *Sanguibacter, Micromonospora, Oerskovia* and *Saccharopolyspora*. Genomic analysis of the strains revealed a large phylogenetic distance to currently known strains and much uncharacterised biosynthetic potential. Obtaining multiple replicates from mammoths is not feasible, and minimizing the contamination risk during the excavation process is therefore particularly important. The specimen was transported and stored in a frozen condition. As the intestines that we encountered were still intact, we could open them under sterile conditions and extract faeces with a low risk of external contamination. Extreme caution was taken to carefully remove the faecal sample, and the specific sample we eventually used for isolation of the Actinobacteria was then extracted from the very core of the large faecal sample, so as to eliminate the risk of contamination to the absolute minimum. The intestinal lumen was thoroughly inspected for defects, and faeces was taken and stored using sterile materials.

A previous metagenome-based study revealed the presence of actinobacterial reads in different tissues of mammoth remains ^35^. However, studying metagenome data always has the risk for misassembly of (damaged) DNA, which is not the case when isolates are grown up and then sequenced. In addition, ancient DNA is more difficult to isolate since it is often fragmented and available in limited amounts ^36^. This is also reflected in the metagenome we obtained, where only low concentrations of DNA could be isolated. As a result, only a fraction of the complete genomic diversity could be analysed. Therefore, in this study, we isolated Actinobacteria from such an environment, strengthening previous findings and allowing us to further explore their functional role and potential. Taxonomic analysis revealed a significant phylogenetic difference between the isolates and their closest known neighbours, strongly suggesting that these isolates represent novel species. The low similarity to known strains may be attributed to the limited number of publicly available complete genomes for underexplored taxa such as *Oerskovia, Sanguibacter* and *Saccharopolyspora*, with three, four and eight available complete genome sequences, respectively, as compared to 323 complete *Streptomyces* genomes. Nevertheless, our *Streptomyces* isolates also showed low similarity scores, while many publicly available complete genomes were available for comparison. The mammoth isolates are therefore a valuable addition to the database and illustrate the power of harvesting microbes from unique environments, even for well-studied genera.

Ancient samples allow a glimpse into the past and may provide evolutionary insights. An interesting question that arises is what the significance is of the major differences that we observe between the isolates and their closest known relatives. 28,500 years on a scale of 2,700 million years reflects only a fraction of actinobacterial evolution, but theoretically, 10-20% ANI variation could be obtained within this time frame. However, the phylogenetic distance between the isolates and their known relatives makes it hard to evaluate evolutionary differences on a genomic level. For better comparison, genome sequences of more closely related strains are required; however, it is yet not known if such strains can be found in the available strain collections.

Much interest has been directed towards the presence of antibiotic-producing Actinobacteria in the mammalian microbiome as this could point towards a protective role against infection by pathogens ^37, 38^. We were excited to be able to isolate six Actinobacteria from the faecal sample. While we do appreciate that this may seem like a low number, it is unknown how well spores survive such a long period of time. Furthermore, Actinobacteria represent only a very minor fraction of the mammalian gut, in particular when compared to soil samples; still, antibiotic-producing Actinobacteria have been isolated from the faeces of a variety of mammals ^39-41^. Studies analysing the microbiome composition of elephants, the closest living relative to mammoths, report the presence of actinobacterial reads in low abundance, including *Streptomyces, Sanguibacter, Micromonospora* and *Saccharopolyspora*, but not *Oerskovia* ^42-44^. Species of *Streptomyces* and *Micromonospora* have been isolated from the faeces of elephants ^41, 45^. However, these isolates have not been sequenced and could not be compared to the bacteria isolated in this study. Alternatively, the isolated Actinobacteria may not have been commensal to the mammoth gut, but instead may have been obtained as part of digested plant material.

Genome mining of the isolates revealed a wide variety of BGCs, with 19% encoding PKSs, 14% NRPSs and 14% terpene biosynthetic enzymes. Most of the predicted BGCs showed low or no similarity to any known BGCs. Comparative genome analyses between M19 and its taxonomic neighbours found in publicly available sequence data revealed a shared conserved internal region and a less conserved region near the ends of the chromosome, holding high numbers of unique and uncharacterised BGCs. This is consistent with results from previous studies showing that common BGCs are often located in the internal regions of the chromosome, while more unique genes are located towards the subtelomeric regions ^9, 46, 47^. Surprisingly, only two BGCs, encoding the desferrioxamine and ectoine pathways, were shared between isolate *Streptomyces* sp. M19 and its neighbours. These specialised metabolites play an important role in survival and their BGCs are well conserved among Actinobacteria^9, 48^. The limited amount of shared biosynthetic content between *Streptomyces* sp. M19 and its neighbours is unexpected as phylogeny has been shown to be an important indicator of BGC distribution ^49, 50^. This underlines the opportunities offered in terms of the biosynthetic potential of bacteria isolated from underexplored environments.

The majority of the isolates displayed bioactivity against one of the tested indicator strains, both through production of volatile and non-volatile compounds. The antibacterial activity against Gram-positive bacteria by *Oerskovia* sp. M15 is surprising, because to the best of our knowledge, antibacterial activity has never been reported before for members of this genus. Four potential BGCs were identified in the genome sequence of M15, of which three had similarity scores below 12%. However, the crude extracts did not show any activity and chemical dereplication did not give any hits with known bioactive compounds. We aim to identify the BGC linked to the bioactivity of *Oerskovia* sp. M15 and identify its cognate natural product(s). The crude extracts of *Streptomyces* isolates M10 and M19 exhibited antibiotic activity against *B. subtilis*. Molecular networking and chemical dereplication using the GNPS platform resulted in the annotation of several known natural products, including antimycin and monactin in the extracts of M10, and conglobatin in the extracts of M19. This was supported by the detection of their BGCs by antiSMASH. Interestingly, antimycin- and conglobatin-related molecules were also detected in the crude extracts of polar Actinobacteria isolated from ancient sediment cores ^12^. An evolutionary path of the antimycin BGC has been proposed in which the L-form BGC is appointed as the ancestor of other antimycin BGCs ^51^. Therefore, it’s not surprising that our detected ancient BGC is the ancestral L-form antimycin cluster. Additionally, analysis of the biogeographical and phylogenetic distribution of the antimycin BGC has revealed that this BGC is widespread among *Streptomyces spp*. and across the globe ^51^. While these known metabolites might be responsible for the bioactivity of the crude extracts, other unknown metabolites with antibacterial activity might be produced as well. The molecular network of the metabolome of these isolates revealed several spectral families unique to the isolates which could not be dereplicated, which may represent novel chemistry, especially for the underexplored genera *Micromonospora, Saccharopolyspora, and Oerskovia*.

In summary, we have isolated Actinobacteria from a unique ancient mammoth stool sample. The large phylogenetic distance between the isolates and their modern siblings as well as the high percentage of uncharacterised biosynthetic potential in the ancient Actinobacteria, even in the representatives of the well-studied genus *Streptomyces*, illustrate that we have by no means captured all microbial diversity. Future studies on the evolutionary differences between the isolates and their modern siblings may provide a unique glance into microbial history.

## Materials and Methods

### Sample acquisition

In February 2014 the specimen was transported by truck to Yakutsk. Dissection of the specimen was done in the autopsy room of the medical faculty of the university of Yakutsk. The samples for microbiological analysis were extracted from 10-14 March 2014, the timespan in which the remains completely thawed. As thawing set on from the outside of the carcass and slowly proceeded inward, each day freshly thawed samples could be taken. Approached from the right side, the intestines where exposed on 13 March after carefully removing soft-tissue and ribs over the previous days. A 60 cm intestinal specimen was taken out of the remains and placed on a sterile surface for further examination. No defects of the intestinal lumen were found. After inspection, two perpendicular (5 cm) incisions where made, and the intersection was folded over, exposing the intestinal lumen. Faecal samples were carefully taken from the lumen using flocked swab collecting tubes (eSwab Copan) or deposited in sterile collection tubes with disposable tweezers with minimal possible cross-contamination. Sterile examination gloves and instruments were used during the whole procedure.

### Isolation of Actinobacteria

Approximately 100 mg of faeces was aseptically placed into a sterile Eppendorf, dissolved in dH_2_O, and serially diluted (10^−1^–10^−6^). Serial dilutions were plated onto different agar media. The media were glucose agar (GA) ^52^, humic acid agar (HA) ^53^, mannitol soya flour medium (SFM) ^54^, modified Starch-Casein agar (MSCA) ^55^, minimal medium without carbon sources (MM) ^54^, MM + 1% glycerol (w/v) (Y), MM + 1% mannitol (w/v) (A). All media contained nystatin (50 µg/mL) and nalidixic acid (10 µg/mL) for the inhibition of fungi and Gram-negative bacteria respectively. Plates were incubated at 30 °C, 4 °C, and anaerobically at room temperature. Single actinomycete colonies were streaked onto SFM agar plates until pure and cryopreserved with glycerol (20 %) and stored at −80 °C.

### Genome sequencing

Strains were cultured in TSBS at 30 °C with 200 rpm shaking speed. Genomic DNA was isolated by phenol-chloroform extraction as described previously ^54^ and sent to be commercially sequenced at Future Genomics Technologies, The Netherlands. Genomes were sequenced using the MinION Nanopore sequencing platform and Illumina NovaSeq6000. Hybrid assembly (both Illumina and ONT reads) was performed for each isolate using Unicycler (v0.4.0.7) ^56^. Briefly, Unicycler performs a SPAdes assembly of the Illumina reads and then scaffolds the assembly graph using long reads. Unicycler polishes its final assembly with Illumina reads and uses Pilon ^57^ to reduce the rate of small base-level errors. The genomes have been deposited at GenBank under accession numbers JAMYWI000000000 (*Sanguibacter* sp. M9), JAMYWH000000000 (*Streptomyces* sp. M10), JAMYWG000000000 (*Micromonospora* sp. M12), JAMYWF000000000 (*Oerskovia* sp. M15), JAMYWE000000000 (*Streptomyces* sp. M19), and CP098407 (*Saccharopolyspora* sp. M46).

### Metagenome sequencing

Genomic DNA of the ice and faecal sample was isolated and sent to be commercially sequenced at Future Genomics Technologies, The Netherlands. There they quantified the gDNA concentration using a Qubit fluorometer and TapeStation system. Since the amounts of detectable gDNA were low, a QIAGEN REPLI-g kit was applied on the samples (Table S4). This successfully increased the gDNA concentration of the ice sample, whereafter sequencing is performed with the Illumina NovaSeq6000. Sequencing adapters were removed by applying the read trimming tool Trimmomatic v0.39 ^58^.

### Phylogenetic analysis

The initial taxonomy of the six isolates was determined using 16S rRNA sequences and BLASTN, with the highest scoring sequencing hits reported. To determine the phylogenetic class of the isolates on a whole genome scale, 578 Actinobacteria genome sequences have been downloaded from the PhyloPhlAn v3.0 ^59^ database, using the phylophlan_get_reference function. Next, the phylophlan_write_config_file script is employed to create a configuration file with DIAMOND as mapping tool, MAFFT for the multiple sequence alignment, trimAl for alignment trimming and IQ-TREE for generating a phylogenetic tree. FastANI v1.32 ^60^ was used to calculate the relatedness of the isolates and neighbouring strains.

### Bioactive potential and comparative genome analysis

AntiSMASH v6.0 ^30^ was used under default settings to predict BGCs from the six isolated bacteria and their modern-day closest known neighbours. The neighbours of *Streptomyces* sp. M19 were downloaded from NCBI using accession numbers: GCA_000147815.3 (*Streptomyces violaceusniger* Tu 4113), GCA_000418455.1 (*Streptomyces rapamycinicus* NRRL 5491), GCA_002812405.1 (*Streptomyces* sp. M56) and GCA_900105695.1 (*Streptomyces melanosporofaciens*). These strains are used as input for pangenome graph builder tool pggb v0.4.0 ^61^ using 90% sequence identity and a segment length of 10000.

### Stereomicroscopy and Scanning Electron Microscopy (SEM)

Isolates were grown for 9 days on SFM. Stereo microscopy was done using a Leica MZ16 FA microscope equipped with a Leica DFC420 C camera. SEM studies were performed using a JEOL JSM-7600F scanning electron microscope ^62^. Single colonies were excised from agar plates and the bottom layer of agarose was cut off to minimise the thickness of the sample. The sections were glued upon a Cryo-EM stub and the whole stub was submerged in non-boiling liquid nitrogen and frozen for 20 s. The stub was then transferred to the gatan Cryo unit, heated to −90 °C for 2 minutes to remove any ice crystals formed during the transfer, cooled to −120 °C, and coated with gold/palladium (80/20) using a sputter coater. Hereafter, the samples were transferred into the microscope and kept at −120 °C while imaging.

### Antimicrobial activity assays

*Bacillus subtilis* 168, *Escherichia coli* ASD19 ^63^, and *Pseudomonas aeruginosa* PA01 were used as indicator strains for antimicrobial activity and were cultured in LB media at 37 °C ^64^. Antimicrobial activity assays were conducted in liquid and on plate, using different methods: *Liquid-grown cultures*: strains were grown in ISP2 (DSMZ #987) and NMMP ^54^ medium for 7 days. Wells were created in soft LB agar (1.8% w/v agar) containing one of the indicator strains pre-grown in liquid LB to exponential phase (OD_600_ = 0.4 – 0.6) by using the opposite end of pipette tip. The wells were filled with 100 µL culture supernatant. Plates were incubated overnight at 37 °C (± 18 hours) and the following day, the zone of inhibition was determined. *Cross streak method:* each strain was independently inoculated on NA (Difco) and ISP2 agar plates as a single streak in the centre of the plate and incubated for 7 days to allow the strains to grow and produce antibiotics. The plates were then seeded with the indicator strains pre-grown in liquid LB to exponential phase (OD_600_ = 0.4–0.6) by streaking perpendicular to the line of actinobacterial growth and incubated overnight at 37 °C (± 18 hours). The following day, the inhibition distance was determined.

#### Double-layer agar method

strains were manually spotted (2 µL) on minimal medium agar plates (MM) supplemented with 0.5% mannitol and 1% glycerol (w/v) as non-repressing carbon sources, and Czapek Dox plates. After seven days of incubation, plates were overlaid with soft LB agar (1.8% w/v agar) containing one of the indicator strains pre-grown in liquid LB to exponential phase (OD_600_ = 0.4 – 0.6) and incubated overnight at 37 °C (± 18 hours). The following day, the zone of inhibition was determined.

#### Volatile assay

the antimicrobial activity of volatile compounds was assessed using a petri dish with two equally sized compartments separated by a plastic divider, both filled with NA. Mammoth isolates were streaked on one side and plates were incubated for 7 days, after which *E. coli* or *B. subtilis* were inoculated on the other side at a concentration of 10^4^ and 10^3^ CFU/mL, respectively.

### Metabolite profiling, MS/MS-based molecular networking, and dereplication

Isolates M10, M15, M19 and M46 were grown as confluent lawns on NA plates for seven days. The agar plates were cut into small pieces, soaked overnight in ethyl acetate (EtOAc) to extract the metabolites, evaporated at room temperature and dissolved in methanol (MeOH) to a concentration of 1 mg/mL. LC-MS/MS acquisition and MS/MS based molecular networking was performed as described previously ^65^. The molecular networking job in GNPS can be accessed at https://gnps.ucsd.edu/ProteoSAFe/status.jsp?task=e160b564fc7e48e6b82394991bfd79be.

### Bioactivity crude extracts

The activity of the crude extracts was determined in triplicate. Indicator strains were pre-grown in liquid LB to exponential phase (OD_600_ = 0.4 – 0.6). Cultures were diluted to OD_600_ = 0.01 in LB and 100 µL diluted culture was loaded in wells of a 100-well honeycomb plate. 20 µg crude extract was added to the wells. Additionally, the following controls were added: LB, bacterial dilution (growth control), bacterial cells + 6 µg ampicillin (positive control), and bacterial cells + MeOH (negative control). Subsequently, the OD_600_ was measured every 30 min for 16 hours using a Bioscreen C Reader (Thermo Scientific, Breda, The Netherlands), with continuous shaking. The OD_600_ was plotted against the time.

## Acknowledgements

The authors acknowledge the support from their respective research institutes. The work was supported by the profile area Antibiotics at Leiden University to G.P.v.W. and by the European Union via ERC Starting Grant 948770-DECIPHER to M.H.M.

## Competing interests

The authors declare no competing interests.

## Notes

### Competing Interest Statement

The authors have declared no competing interest.

