## Supplemental Tables and Figures for "Taxonomic and metabolic diversity of Actinobacteria isolated from faeces of a 28,000-year-old mammoth"

**Table S1. Isolate taxonomic classification and identified closest neighbours.**

| Isolate ID | Taxonomic classification | Closest known neighbor(s) | Accession number | ANI score % |
|---|---|---|---|---|
| M9 | *Sanguibacter* | *Sanguibacter antarcticus* | GCA_002564005.1 | 84.0 |
| M10 | *Streptomyces* | *Streptomyces* sp. NTK 937 | GCA_000698495.1 | 84.9 |
| | | *Streptomyces* sp. SM18 | GCA_002910775.2 | 84.8 |
| M12 | *Micromonospora* | *Micromonospora chokoriensis* | GCA_900091505.1 | 89.2 |
| M15 | *Oerskovia* | *Oerskovia enterophila* | GCA_001692445.1 | 88.4 |
| | | *Oerskovia* sp. Root22 | GCA_001429135.1 | 88.4 |
| | | *Oerskovia* sp. Root918 | GCA_001428945.1 | 88.4 |
| M19 | *Streptomyces* | *Streptomyces* sp. M56 | GCA_002812405.1 | 82.3 |
| | | *Streptomyces rapamycinicus* NRRL 5491 | GCA_000418455.1 | 82.4 |
| | | *Streptomyces melanosporofaciens* | GCA_900105695.1 | 82.5 |
| | | *Streptomyces violaceusniger* Tu 4113 | GCA_000147815.3 | 82.3 |
| | | *Streptomyces* sp. MnatMP-M27 | GCA_900092015.1 | 82.1 |
| | | *Streptomyces antioxidans* | GCA_000968685.2 | 82.6 |
| | | *Streptomyces sparsogenes* | GCA_001969965.1 | 82.7 |
| M46 | *Saccharopolyspora* | *Saccharopolyspora flava* | GCA_900116135.1 | 88.0 |

**Table S2. Isolates from extreme environments and their similarity scores with closest neighbours.**

| Isolate ID | Origin | Accession number | Closest known neighbor(s) | Origin neighbour(s) | Accession number neighbour(s) | ANI score % |
|---|---|---|---|---|---|---|
| *Streptomyces* sp. NTK 937 | Deep-sea, Canary Basin, Canary Islands | GCA_000698495.1 | *Streptomyces* sp. SM18 | Deep-sea, Kilkieran Bay, Galway, Ireland | GCA_002910775.2 | 96.7 |
| *Streptomyces* sp. SCSIO 3032 | Deep-sea, Madeira archipelago, Portugal | GCA_002128305.1 | *Streptomyces* sp. MP131-18 | Deep-sea, Trondheim, Norway | GCA_001984575.1 | 85.5 |
| *Streptomyces* sp. BF-3 | Great Salt Plains, Oklahoma | GCA_002104865.1 | *Streptomyces* sp. Cmuel-A718b | Unknown | GCA_900092005.1 | 98.9 |
| *Streptomyces* sp. 4F | Great Salt Plains, Oklahoma | GCA_002104855.1 | *Streptomyces* sp. Alain-F2R5 | Al Ain, Dubai, United Arab Emirates | GCA_002277855.1 | 93.1 |
| ***Streptomyces thermolilacinus* SPC6** | Sand, Linze desert, China | GCA_000478605.2 | *Streptomyces rubrolavendulae* MJM4426 | Soil, Jeju, South Korea | GCA_001750785.1 | 88.2 |

**Table S3. Percentage of BGCs that share less than 50% KnownClusterBlast similarity to BGCs within the MiBIG database of the mammoth isolates and a random selection of Actinobacteria**

| Strain | Isolation source | total # BGCs | % BGC < 50% similarity* |
|---|---|---|---|
| ***Sanguibacter* sp. M9** | mammoth faeces | 6 | 67% |
| *Sanguibacter gelidistatuariae* ISLP-3 | ice sculpture, Antarctica | 3 | 67% |
| *Sanguibacter keddieii* DSM 10542 | bovine blood | 5 | 80% |
| *Sanguibacter massiliensis* | human stool | 2 | 50% |
| ***Micromonospora* sp. M12** | mammoth faeces | 28 | 79% |
| *Micromonospora auratinigra* DSM 44815 | soil, Thailand | 16 | 88% |
| *Micromonospora chokoriensis* DSM 45160 | sandy soil | 15 | 73% |
| *Micromonospora inosilota* DSM 43819 | forest soil | 9 | 89% |
| *Micromonospora narathiwatensis* DSM 45248 | peat swamp forest soil | 18 | 67% |
| *Micromonospora terminaliae* DSM 101760 | plant | 13 | 77% |
| *Micromonospora zamorensis* DSM 45600 | rhizosphere | 12 | 83% |
| ***Oerskovia* sp. M15** | mammoth faeces | 4 | 75% |
| *Oerskovia* sp. KBS0722 | soil | 6 | 83% |
| *Oerskovia* sp. Root22 | roots of *Arabidopsis thaliana* | 3 | 33% |
| ***Saccharopolyspora* sp. M46** | mammoth faeces | 20 | 80% |
| *Saccharopolyspora coralli* E2A | stony coral | 10 | 70% |
| *Saccharopolyspora erythraea* NRRL 2338 | soil | 36 | 72% |
| *Saccharopolyspora* sp. ASAGF58 | soil | 29 | 69% |
| ***Streptomyces* sp. M10** | mammoth faeces | 31 | 65% |
| ***Streptomyces* sp. M19** | mammoth faeces | 34 | 65% |
| *Streptomyces antibioticus* DSM 40234 | soil | 26 | 50% |
| *Streptomyces sp.* MP113-05 | sponge | 28 | 79% |
| *Streptomyces cacaoi* H2S5 | moss soil | 31 | 45% |
| *Streptomyces coelicolor* CFB_NBC_0001 | soil | 27 | 41% |
| *streptomyces ficellus* NRRL 8067 | soil | 26 | 73% |
| *streptomyces galilaeus* ATCC 14969 | soil | 24 | 50% |
| *streptomyces griseus* ATCC 13273 | soil, Japan | 30 | 60% |
| *Streptomyces leeuwenhoekii* C34 | desert soil | 35 | 69% |
| *Streptomyces lincolnensis* NRRL 2936 | soil | 29 | 59% |
| *Streptomyces lunaelactis* MM109 | moonmilk speleothem | 28 | 71% |
| *streptomyces olivochromogenes* DSM 40451 | soil | 39 | 72% |
| *streptomyces scabiei* 87.22 | plant pathogen | 34 | 59% |
| *streptomyces tsukubensis* AT3 | forest soil, China | 36 | 64% |

**Table S4. DNA concentration statistics**

| Sample | Qubit | Tapestation | | | Repli-G |
|---|---|---|---|---|---|
| | Conc. ng/µl | Conc. ng/µl | DIN | peak | Conc. ng/µl |
| **Ice** | 23.20 | 10.00 | 4.3 | 6847 | 218 |
| **Faeces** | 17.20 | 9.90 | 5.4 | 6477 | - |

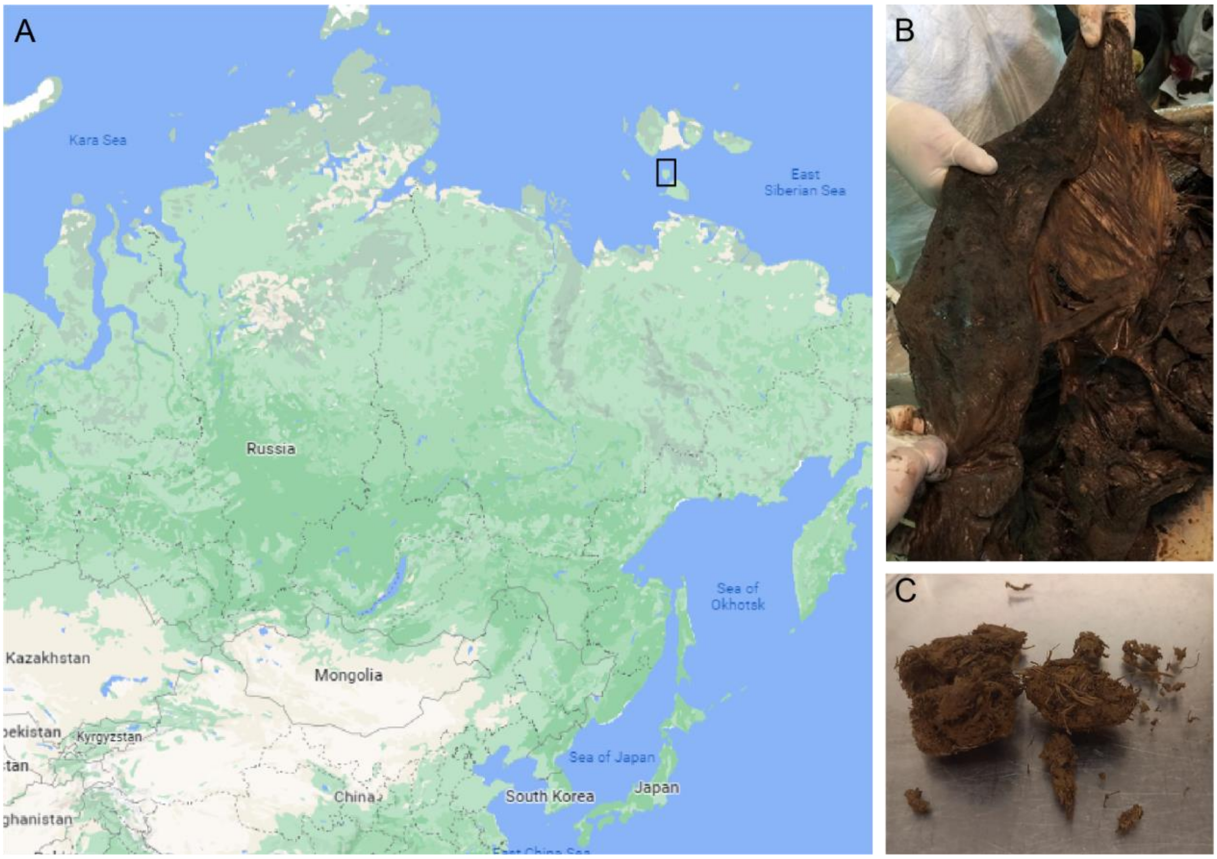


**Figure S1. Origin and visualization of the mammoth sample.** A) the mammoth carcass was discovered in August 2012 on Maly Lyakhovsky Island, Russia. B) extracted 60 cm intestinal specimen without any defects of the intestinal lumen C) faecal sample.

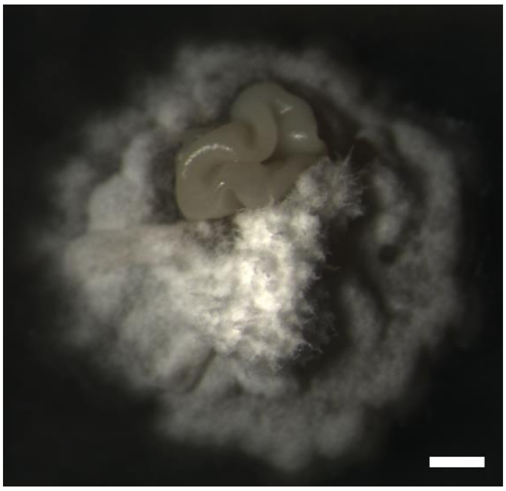

**Figure S2. Stereomicrograph showing morphological heterogeneity within colonies of *Streptomyces* isolate M19.** *Streptomyces* sp. M19 was grown for 12 days on MYM agar. Scale bar, 1 mm.

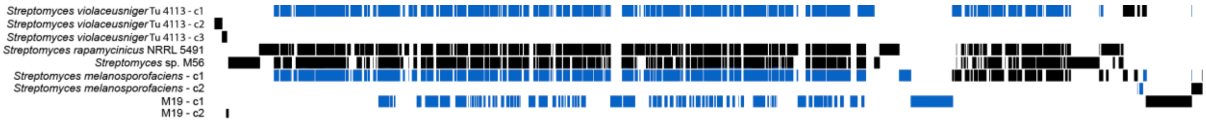


**Figure S3. Pangenome graph of *Streptomyces* sp. M19 and its closest neighbours, *Streptomyces violaceusniger* Tu 4113, *Streptomyces rapamycinicus* NRRL 5491 and *Streptomyces* sp. M56.** Coloured blocks indicate shared areas between isolates. Blue lines indicate reverse orientation.

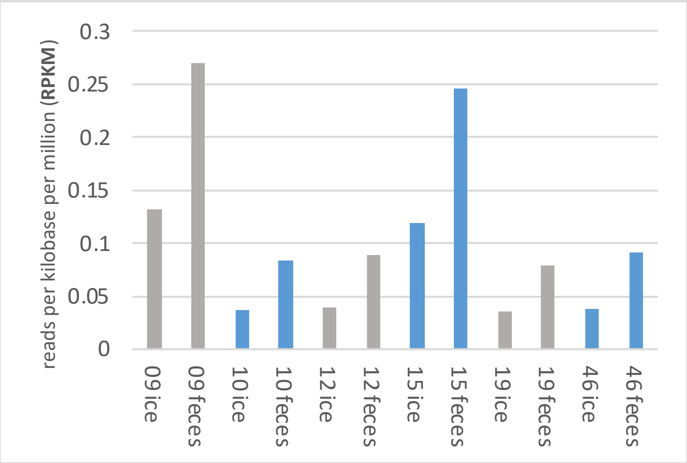


**Figure S4. Metagenomic reads of ice and faeces samples mapped to the isolates.** Normalised read counts for ice and faeces samples are displayed for each of the six mammoth isolates, M9, M10, M12, M15, M19 and M46.

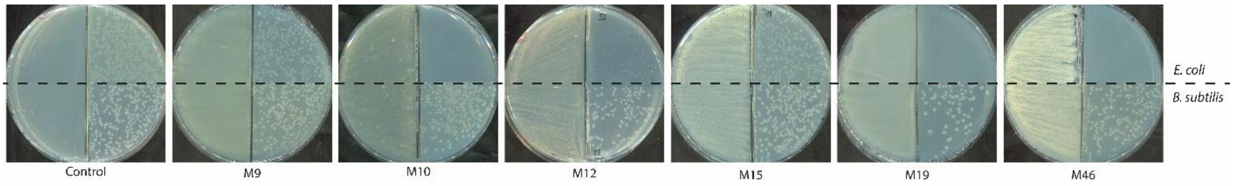


**Figure S5. Bioactivity of volatiles released by the mammoth isolates.** Mammoth isolates were grown for 7 days on NA agar. After this period, indicator strains *E. coli* strain ASD19 and *B. subtilis* 168 were plated on the other side, also on NA agar, and growth was assessed visually. Note that isolates M10, M12, M19, and M46 inhibit growth of *E. coli*.

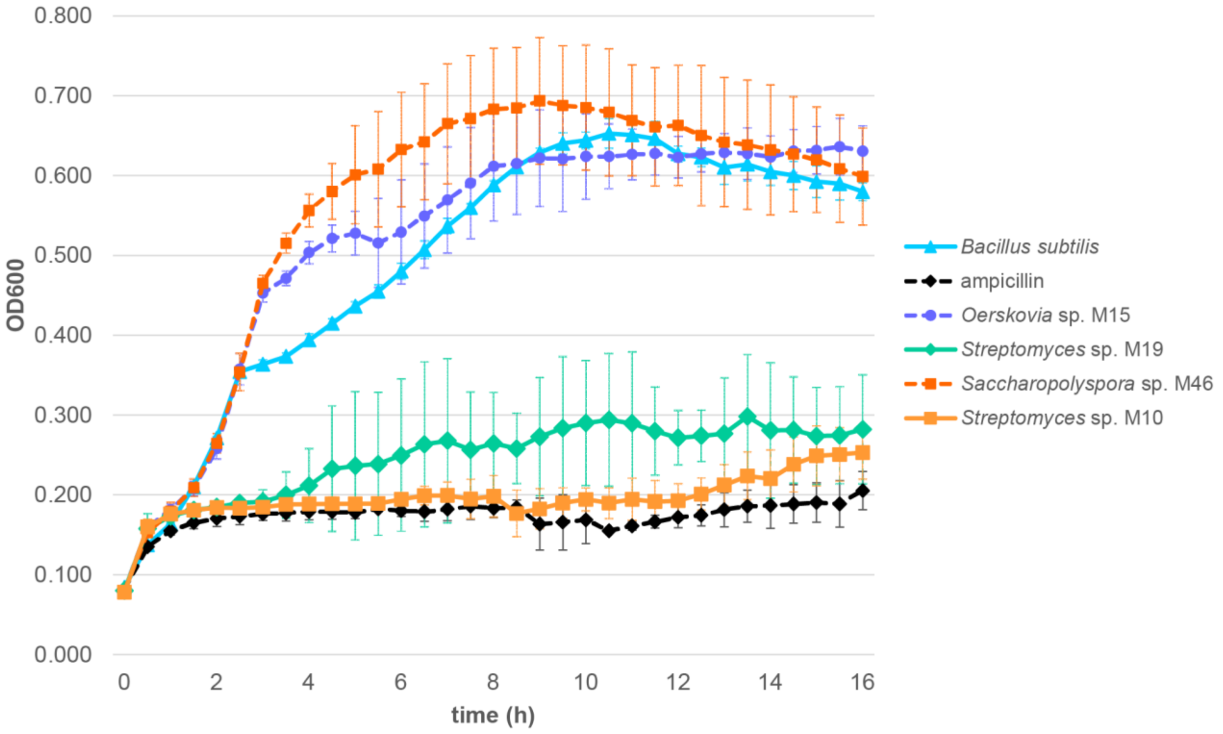


**Figure S6. The crude extracts of *Streptomyces* sp. M10 and M19 inhibit the growth of *B. subtilis.*** To determine the antimicrobial activity of the crude extracts, 20 µg extract was added to 100 µL diluted *B. subtilis* culture (*n* = 3). The $OD_{600}$ was measured every 30 min for 16 hours, after which the $OD_{600}$ was plotted against the time and growth curves were compared to the growth of *B. subtilis* alone. 6 µg ampicillin was added as a positive control.

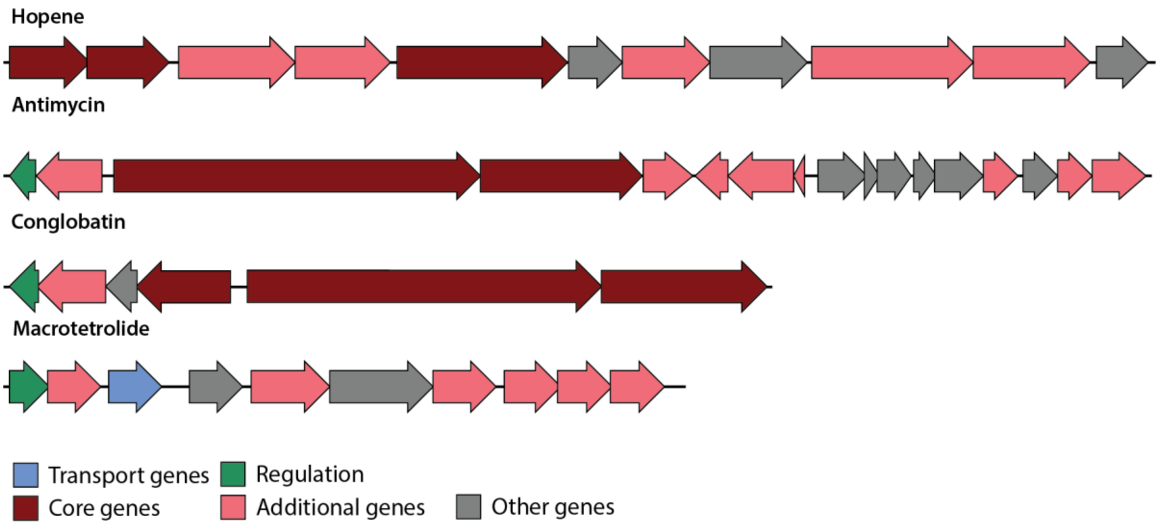


**Figure S7. Predicted hopene cluster of M10 & M19, antimycin and macrotetrolide gene cluster of M10 and the conglobatin gene cluster of M19.** Colours of the genes are based on the colouring system of antiSMASH v6.0.
